# New insights on adaptation and population structure of cork oak using genotyping by sequencing

**DOI:** 10.1101/263160

**Authors:** F. Pina-Martins, J. Baptista, G. Pappas, O. S. Paulo

## Abstract

Species respond to global climatic changes in a local context. Understanding this process is paramount due to the pace of these changes. Tree species are particularly interesting to study in this regard due to their long generation times, sedentarism, and ecological and economic importance. *Quercus suber* L. is an evergreen forest tree species of the Fagaceae family with an essentially Western Mediterranean distribution. Despite frequent assessments of the species’ evolutionary history, large-scale genetic studies have mostly relied on plastidial markers, whereas nuclear markers have been used on studies with locally focused sampling strategies. The potential response of *Q. suber* to global climatic changes has also been studied, under ecological modelling. In this work, “Genotyping by Sequencing” (GBS) is used to derive 2,547 SNP markers to assess the species’ evolutionary history from a nuclear DNA perspective, gain insights on how local adaptation may be shaping the species’ genetic background, and to forecast how *Q. suber* may respond to global climatic changes from a genetic perspective. Results reveal an essentially unstructured species, where a balance between gene flow and local adaptation keeps the species’ gene pool somewhat homogeneous across its distribution, but at the same time allows variation clines for the individuals to cope with local conditions. “Risk of Non-Adaptedness” (RONA) analyses, suggest that for the considered variables and most sampled locations, the cork oak does not require large shifts in allele frequencies to survive the predicted climatic changes. However, more research is required to integrate these results with those of ecological modelling.

## 1 Introduction

### 1.1 Adaptation

Global climatic changes have been shown to cause alterations in species’ traits (Benito Garzón, Alía, Robson, & Zavala, 2011; Walther et al., 2002). Understanding how species respond to such alterations in their environmental context is becoming an increasingly important question due to the pace at which they are taking place (Kremer et al., 2012; Primack et al., 2009). To avoid obliteration, species may respond to climatic changes by either altering their distribution range, effectively going extinct in the original location but persisting somewhere else, or by adapting to the new conditions. The latter can occur “instantly”, due to phenotypic plasticity, or across several generations, by local adaptation (Aitken, Yeaman, Holliday, Wang, & Curtis-McLane, 2008). The kind of response species can provide is known to depend on factors like location, distribution range, and/or genetic background (Gienapp, Teplitsky, Alho, Mills, & Merilä, 2008; Ohlemuller, Gritti, Sykes, & Thomas, 2006).

Tree species are characterized by sedentarism, long lifespan and generation times, allied with generally large distribution ranges and capacity for long distance dispersal through pollen and seeds (Kremer et al., 2012). These traits make them interesting subjects to study regarding their response to global climatic changes (Thuiller et al., 2008).

In this work, we address the case of the cork oak *(Quercus suber* L.). With a distribution ranging most of the West Mediterranean region (Figure 1), this oak species is the most selective evergreen oak of the Mediterranean basin in terms of precipitation and temperature conditions (Vessella, López-Tirado, Simeone, Schirone, & Hidalgo, 2017). European oaks in particular, are known to have endured past climatic alterations, but how they can cope with the current, rapidly occurring changes is not yet fully understood (Kremer et al., 2012; Kremer, Potts, & Delzon, 2014). Despite this tree’s ecological and economic importance, little is known regarding the consequences of global climatic change on its future (Benito Garzón, Sánchez de Dios, & Sainz Ollero, 2008).

**Figure 1:**
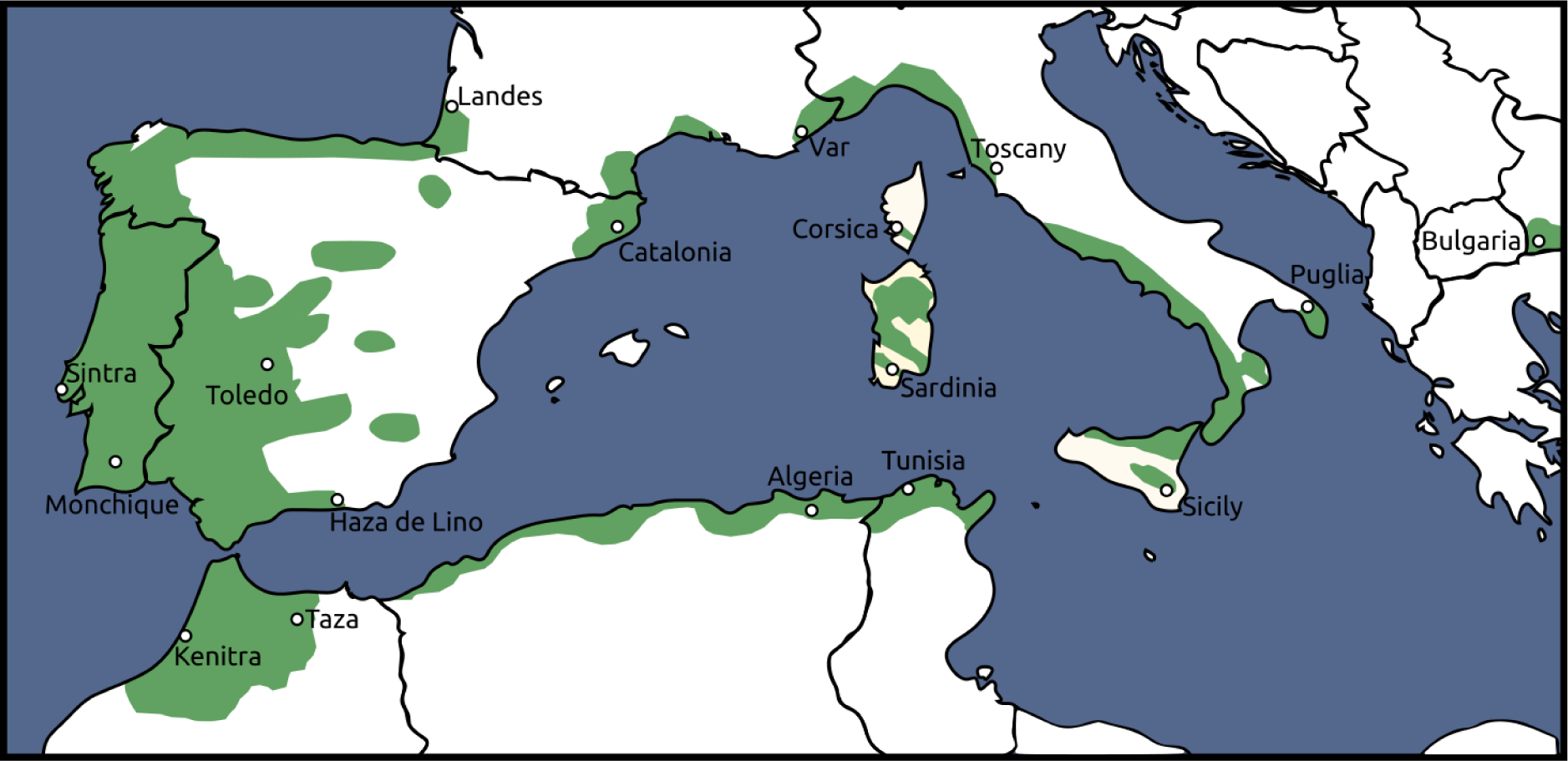
A map of cork oak *(Quercus suber)* distribution. Land areas in green represent the species’ range. White dots represent the sampling locations. Adapted from EUFORGEN 2009 (www.euforgen.org).

Some recent works have been performed to attempt to answer this very question, but focusing on range expansion and contraction with the assumption of a genetically homogeneous species and niche conservationism (Correia, Bugalho, Franco, & Palmeirim, 2017; Vessella et al., 2017). Both these studies also highlight the need for a genetic study regarding the adaptation potential of *Q. suber*. However, studies integrating genetic information and response to climatic alterations of *Q. suber* are rare and of small scale (Modesto et al., 2014) when compared with other oak species (Rellstab et al., 2016). Studies such as (Jose Alberto Ramírez-Valiente, Valladares, Huertas, Granados, & Aranda, 2011) have revealed that some traits can be associated to genetic variants, however, these were performed on a local scope and using a relatively low number of markers, which limits their utility in a larger scope. Knowing gene flow and local adaptation dynamics of *Q. suber*is paramount to understanding the species’ potential to endure rapid climatic changes through adaptation (Savolainen, Lascoux, & Merilä, 2013).

Genomic resources represent a new way to study the genetic mechanisms responsible for local adaptation (Rellstab, Gugerli, Eckert, Hancock, & Holderegger, 2015), through the use of environmental association analyses, which correlate environmental data with genetic markers, thus highlighting loci putatively involved in the adaptation process (Rellstab et al., 2016). The same methods, can thus, in principle, be used to assess the degree of maladaptation to predicted future local conditions (Rellstab et al., 2016). Applying this kind of methodology on *Q. suber* would fill the gap mentioned in (Correia et al., 2017; Vessella et al., 2017), that multidisciplinary approaches are required to more accurately provide sound recommendations for the conservation of forests. The Risk of Non-Adaptedness (RONA) method was developed in (Rellstab et al., 2016) with this very goal, however, no public implementation is provided in the mentioned work.

### 1.2 Population structure

In order to predict a species’ response to change (Kremer et al., 2012), it is fundamental to know both its genetic architecture of adaptive traits (Alberto et al., 2013) and evolutionary history (Kremer et al., 2014). However, the very nature of genetic and genomic data hampers the distinction of selection signals from other processes (McVean & Spencer, 2006), especially demographic events (Bazin, Dawson, & Beaumont, 2010). In order to overcome the obstacles caused by the entanglement of population structure (mostly shaped by gene flow, inbreeding, and genetic drift) and selection (Foll, Gaggiotti, Daub, Vatsiou, & Excofer, 2014), recent methods incorporate population structure information to detect adaptation (Gautier, 2015; Günther & Coop, 2013). Likewise, methods to accurately estimate population structure should be performed without loci known to be under selection (De Kort et al., 2014).

The evolutionary history of *Q. suber* has been studied in the past using multiple methodologies and in different geographic ranges. The most recent large-scale studies on the subject suggest that cork oak is divided into four strictly defined lineages (Magri et al., 2007; Simeone et al., 2009). Two of these lineages range from the south-east of France, to Morocco, including the Iberian peninsula and the Balearic Islands, a third lineage ranges from the Monaco region to Algeria and Tunisia, including the islands of Corsica and Sardinia. The fourth lineage spans the entire Italic peninsula, including Sicilia. Based only on plastidial markers, these lineages have been shown to hardly share any haplotypes. Notwithstanding, later works based on nuclear DNA have hinted at a different scenario, where the species is not as strictly divided (Costa et al., 2011; J. A. Ramírez-Valiente, Valladares, & Aranda, 2014). These works are, however, limited in either geographic scope or number of markers to confidently conclude that such segregation is only present in plastidial markers.

In the present work, a panel of Single Nucleotide Polymorphism (SNP) markers derived from the Genotyping by Sequencing (GBS) technique (Elshire et al., 2011) was developed to attain the following goals: (1) attempt to infer the species’ genetic structure and evolutionary history, (2) detect signatures of natural selection, and (3) investigate the adaptation potential of *Q. suber* based on the RONA method developed and presented on (Rellstab et al., 2016).

## 2 Material & Methods

### 2.1 Sample and environmental data collection

In order to provide a comprehensive view of the species genetic background, samples were collected from 17 locations spanning most of *Q. suber’s* distribution. Fresh leaves were collected from six individuals from, *Bulgaria, Corsica, Kenitra, Monchique, Puglia, Sardinia, Sicilia, Tuscany, Tunisia* and *Var*, and from five individuals from *Algeria, Catalonia, Haza de Lino, Landes, Sintra, Taza* and *Toledo* for a total of 95 individuals (Table 1, Figure 1). It is worth noting that trees from Bulgaria are not of natural origin, but rather the result of human introduction from Iberian locations (Borelli & Varela, 2000; Petrov & Genov, 2004).

**Table 1:** Coordinates and number of sampled individuals for every sampling site.

| Sample site | Latitude (decimal deg.) | Longitude (decimal deg.) | Number of sampled individuals |
| --- | --- | --- | --- |
| Algeria | 36.5400 | 7.1500 | 5 |
| Bulgaria | 41.43 | 23.17 | 6 |
| Catalonia | 41.8500 | 2.5333 | 5 |
| Corsica | 41.6167 | 8.9667 | 6 |
| Haza de Lino | 36.8333 | -3.3000 | 5 |
| Kenitra | 34.0833 | -6.5833 | 6 |
| Landes | 43.7500 | -1.3333 | 5 |
| Monchique | 37.3167 | -8.5667 | 6 |
| Puglia | 40.5667 | 17.6667 | 6 |
| Sardinia | 39.0833 | 8.8500 | 6 |
| Sicilia | 37.1167 | 14.5000 | 6 |
| Sintra | 38.7500 | -9.4167 | 5 |
| Taza | 34.2000 | -4.2500 | 5 |
| Toledo | 39.3667 | -5.3500 | 5 |
| Tunisia | 36.9500 | 8.8500 | 6 |
| Tuscany | 42.4167 | 11.9500 | 6 |
| Var | 43.1333 | 6.2500 | 6 |
| **Total:** | **-** | **-** | **95** |

Most samples were collected from an international provenance trial (FAIR I CT 95 0202) established at “Monte Fava”, Alentejo, Portugal (38°00’ N; 8°7’ W) (Varela, 2000), except Portuguese and Bulgarian samples, which were collected directly from their native locations. The collected plant material was stored at −80°C until DNA extraction.

Altitude, latitude and longitude spatial variables (Varela, 2000) were recorded for each of the native sampling sites. Nineteen Bioclimatic (BIO) variables, BIO1 to BIO19 were collected from the WorldClim database (Hijmans, Cameron, Parra, Jones, & Jarvis, 2005) at 30 arc-seconds (~ 1 km) resolution for “Current conditions ~1960-1990” and “Future” predictions for 2070 (using two different *Representative Concentration Pathways* (RCPs), *rcp26* and *rcp85* conditions for the following “Global Climate Models” (GCMs): BCC-CSM1- 1, CCSM4, GFDL-CM3, GISS-E2-R, HadGEM2-ES, IPSL-CM5A-LR, MRI-CGCM3, MPI-ESM-LR and NorESM1-M as these are available under permissive licenses and calculated for both *rcp26* and *rcp85).* An average of the mentioned datasets was obtained for each coordinate and variable used in the analyses (Supplementary Table 1 and 2 respectively). Data was extracted from the GeoTiff files using a python script, *layer_data_extractor.py* (https://github.com/StuntsPT/Misc_GIS_scripts) as of commit “bd36320”.

Correlations between present Bioclimatic variables were assessed using Pearson’s correlation coefficient as implemented in the R script *eliminate_correlated_variables.R* (https://github.com/JulianBaur/R-scripts) as of commit “43e6553”, which resulted in the exclusion of six variables due to high correlation (*r*>0.95). Each sampling location was thus characterized by three spatial variables and 13 environmental variables (Supplementary Table 3).

### 2.2 Library preparation and sequencing

Genomic DNA was extracted from liquid nitrogen grounded leaves of all samples collected for this work using the kit “innuPREP Plant DNA Kit” (Analytik Jena AG), according to the manufacturer’s protocol.

The total amount of extracted DNA was quantified by spectrophotometry using a Nanodrop 1000 (Thermo Scientific) and integrity verified on Agarose gel (0.8%). DNA samples were then diluted to a concentration of ~100 ng/µl and plated for genotyping. DNA samples were then outsourced to “Genomic Diversity Facility”, at Cornell University” for genotyping using the “Genotyping by sequencing” (GBS) technique as described in (Elshire et al., 2011). Samples were shipped in a single 96 well plate with one “blank” well for negative control. Sequencing was performed according to the standard protocol on a single Illumina HiSeq 2000 flowcell using the low frequency cutter enzyme “EcoT22I”, due to the large size of *Q. suber’s* genome.

### 2.3 Genomic data analyses

The raw GBS data was analysed using the program *ipyrad* v0.5.15, which is based on *pyrad* (Eaton, 2014), using the provided *“condal’* environment - *MUSCLE* v3.8.31 (Edgar, 2004) and *>VSEARCH*> v2.0.3 (Rognes, Flouri, Nichols, Quince, & Mahé, 2016). Sequence assembly was performed for the GBS *datatype*, with the parameters *clustering threshold* of 0.95, *mindepth* of 8 and maximum *barcode* mismatch of 0. Each sampling site had to be represented by at least three individuals for a SNP to be called, except the locations of *Kenitra* and *Taza*, where only one individual was required, due to the lower representation of these sampling sites. Full parameters can be found in Supplementary Datafile 1. The demultiplexed “fastq” files were submitted to NCBI’s Sequence Read Archive SRA) as “Bioproject” PRJNA413625.

Downstream analyses were automated using *“GNU Make”.* This file, containing every detail of every step of the analyses for easier reproducibility is presented as Supplementary Datafile 2. For improved reproducibility, a docker image with all the software, configuration files, parameters and the *Makefile*, ready to use is also provided (https://hub.docker.com/r/stunts/q.suber_gbs_data_analyses/). The intent is not to allow the analyses process to be treated as a “black box”, but rather to provide a full environment that can be reproduced, studied and modified by the scientific community.

Processed data from *ipyrad* was then filtered using *VCFtools* v0.1.14 (Danecek et al., with the following criteria: each sample has to be represented in at least 55% of the SNPs, and after this each SNP has to be represented in at least 80% of the individuals. Furthermore, due to the relatively small sample size, the minimum allele frequency (MAF) of each SNP has to be at least 0.05 for it to be retained.

In order to minimize the effects of linkage disequilibrium, downstream analyses were performed using only one SNP per locus, by discarding all but the SNP closest to the centre of the sequence in each locus. This sub dataset was obtained using the python script *vcf_parser.py* (https://github.com/CoBiG2/RAD_Tools/blob/master/vcf_parser.py) as of commit “0893296”.

All file format conversions were performed using *PGDSpider*v2.1.0.0 (Lischer & Excoffier, 2011), except for the *BayPass* and *SelEstim* formats, where the scripts *geste2baypass.py* (https://github.com/CoBiG2/RAD_Tools/blob/master/geste2baypass.py) and *gest2selestim.sh* (https://github.com/Telpidus/omics_tools) as of commit “b99636e” and “f74f66b” respectively were used, since *PGDSpider* did not handle either of these formats at the time of writing.

Descriptive statistics, such as Hardy-Weinberg Equilibrium (HWE), F_ST_ and F_IS_ were calculated using *Genepop* v4.6 (Rousset, 2008). The same software was further used to perform Mantel tests to determine an eventual effect of Isolation by Distance (IBD) by correlating “‘F/(1-F)’-like with common denominator” with “Ln(distance)” following on 1,000,000 permutations. This test was performed excluding individuals sampled from *Bulgaria* due to their introduced origin.

### 2.4 Outlier detection and environmental associations

Outlier detection was performed using two programs: *SelEstim* v1.1.4 (Vitalis, Gautier, Dawson, & Beaumont, 2014) (50 pilot runs of length 1,000 followed by a main run of length 10^6^, with a burnin of 1,000, a thinning interval of 20, and a detection threshold of 0.01) and *BayeScan* v2.1 (Foll & Gaggiotti, 2008) (20 pilot runs of length 5,000 followed by a main run of 500,000 iterations, a burnin of 50,000, a thinning interval of 10, and a detection threshold of 0.05) (full commands and parameters available in Supplementary Datafile 2), since these methods show the lowest rate of false positives (Narum & Hess, 2011; Vitalis et al., 2014). Only SNPs indicated as outliers by both programs were considered outliers for the purpose of this work. This was done to reduce the chance of false positives, which is a known issue in this type of analyses (Gautier, 2015; Vitalis et al., 2014).

The software *BayPass* v2.1 (Gautier, 2015) wrapped under the script *Baypass_workfow.R* (https://github.com/StuntsPT/pyRona/blob/master/Baypass_workfow.R) as of commit “5b406fb” was used to assess associations of SNPs to environmental variables using the “AUX” model (20 pilot runs of length 1,000, followed by a main run of length 500,000 with a burnin of 5,000 and a thinning interval of 25). Any association with a Bayes Factor (BF) above 15 was considered significant. Similar to what was done for the Mantel tests, association analyses were performed excluding individuals from *Bulgaria* sampling site.

Sequences containing outlier loci or SNPs associated to an environmental variable were queried against the genome of *Q. lobata* (Sork et al., 2016) v1.0 using BLAST v2.2.28+ (Altschul et al., 1997) with an e-value threshold of 0.00001.

### 2.5 Population Structure

Three distinct methods were used for clustering the individuals in order to understand the general pattern of individual or population grouping, namely, Principal Components Analysis (PCA), structure (Pritchard, Stephens, & Donnelly, 2000) and *MavericK* (Verity & Nichols, 2016).

Principal Components Analysis analysis was performed with *snp_pca_static.R* (https://github.com/CoBiG2/RAD_Tools/blob/master/snp_pca_static.R) as of commit “bb2fc45”.

In order to correctly interpret clustering analyses results, it is important to estimate the value of “K”, which represents how many *demes* the data can be clustered into. The structure method was performed with structure v2.3.4, (Pritchard et al., 2000) using the admixture model with an inferred *alpha.* To achieve best results using structure, 20 replicates of each “K” were run at 200,000 iterations (10% burnin), and the three best values of delta K were then run for a single replicate at 2,000,000 iterations (10% *burnin).* The software *MavericK* is especially interesting for cluster estimation due to its innovative method for estimating “K”, called “Thermodynamic Integration” (TI), which has shown superior performance in this task relative to other methods (Verity & Nichols, 2016). In this case, two runs were performed: an initial single “pilot” run of 5,000 iterations, with a *burnin* of 500 using an admixture model, a free *alpha* parameter of “1” and “thermodynamic integration” (TI) turned of. Tuned *alpha* and *alphaPropSD* values were extracted from the pilot run and used in the “tuned” run as parameters for the admixture model. This run was comprised of five runs of 10,000 iterations (10% burnin), with TI turned on and set to 20 rungs of 10,000 samples with 20% burnin. Both programs were wrapped under *Structure_threader* v 1.2.2 (Pina-Martins, Silva, Fino, & Paulo, 2016) for values of “K” between 1 and 8. The most suitable value of “K” was calculated using the *evanno* (Earl & vonHoldt, 2012) and TI methods for and structure and *MavericK* respectively. Full parameter files are available as Supplementary Datafile 2.

In order to obtain an unbiased population structure, the same methodology was used on two more datasets derived from the original data. On one, only SNPs considered outliers or that were associated with environmental variables were used (“non-neutral” dataset), and on the other one, these markers were removed (“neutral” dataset).

### 2.6 Risk of non-adaptedness

The software *pyRona* was developed in this work as the first public implementation of the method described in (Rellstab et al., 2016) called “Risk of non-adaptedness” (RONA). This method provides a way to represent the theoretical average change in allele frequency at loci associated with environmental variables required for any given population to cope with changes in that variable. The program source code is hosted on github, under a GPLv3 license, and can be downloaded free of charge at https://github.com/StuntsPT/pyRona.

In short, for every significant association between a SNP and an environmental variable, the RONA method plots each location’s individuals’ allele frequencies (corrected by *Baypass* to eliminate any possible effects of population structure) vs. the respective environmental variable. This is done for both the current value and the future prediction. A correlation between allele frequencies and the current variable values is then calculated and the corresponding best ft line is inferred. The distance between the fitted line and the two coordinates is then compared per location and its normalized difference is considered the RONA value for each association and location (which can vary between 0 and 1). In theory, the higher the difference in conditions between current values and the prediction, the more *Q. suber* should have to shift its allele frequencies to survive in the location under the new conditions.

Two alternative climate prediction models were used to calculate a RONA value for each location, a low emission scenario (RCP26) and a high emission scenario (RCP85) (IPCC, 2014) in order to account for uncertainties in the models’ assumptions.

The software version 0.1.3 was used and any associations fagged by *Baypass* with a BF above 15 were considered relevant and included in the RONA analysis. Results for the three most frequent non-geospatial environmental variables associated with most SNPs, were selected as the most interesting for determining generic RONA values.

## 3 Results

Genotyping by sequencing (Elshire et al., 2011), a technique based on restriction enzyme genomic complexity reduction followed by short-read sequencing, was employed to discover SNP markers from a total of 95 *Q. suber* individuals sampled from 17 geographical locations (Table 1).

A total of 225,214,094 reads (100 bp) generated by the GBS assay was processed by *ipyrad* (Eaton, 2014) computational pipeline. The first analytical step consisted in the assembly of raw reads into 7,456 distinct contiguous sequence fragments (genomic loci), from which an initial set of 12,330 SNPs were fagged. Twelve *Q. suber* samples were discarded due to low sequence representation during the assembly process, resulting in the retention of 83 individuals. After filtering according to the criteria presented in the methods section 3.3, 2,547 SNPs remained, which were used for all further analyses. This filtering process also further removed two samples due more having more than 55% missing data, and therefore, of the 83 remaining samples, only 81 were used in the analyses (Table 2).

**Table 2:** Number of individuals used in analysis, Fis values, and Hardy-Weinberg Equilibrium (HWE) *p*-values for each sampling site.

| Sample site | Number of individuals used in analysis | $F_{IS}$ | HWE (Het. Def. P-value) | HWE (Het. Exc. P-value) |
| --- | --- | --- | --- | --- |
| Algeria | 4 | 0.09 | **0** | 1 |
| Bulgaria | 4 | 0.01 | 0.26 | 1 |
| Catalonia | 5 | 0.03 | **0** | 1 |
| Corsica | 6 | 0.1 | **0** | 1 |
| Haza de Lino | 5 | 0.04 | **0** | 1 |
| Kenitra | 3 | 0.06 | **0** | 1 |
| Landes | 4 | -0.02 | 0.94 | 0.57 |
| Monchique | 5 | 0.03 | **0** | 1 |
| Puglia | 6 | 0.1 | **0** | 1 |
| Sardinia | 6 | 0.07 | **0** | 1 |
| Sicilia | 3 | 0.09 | **0** | 1 |
| Sintra | 3 | 0.09 | **0** | 1 |
| Taza | 4 | 0.09 | **0** | 1 |
| Toledo | 5 | 0.02 | **0.02** | 1 |
| Tunisia | 6 | 0.05 | **0** | 1 |
| Tuscany | 6 | 0.06 | **0** | 1 |
| Var | 6 | 0.01 | **0.02** | 1 |
| **Total:** | **81 -** |  | **15** | **0** |

The calculated FIS values for each sampling site are available in Table 2. These range from −0.0234 *(Landes)* to 0.0987 *(Puglia)* with an average value of 0.0531. Pairwise FST values are available in Figure 2 and Supplementary Table 4. These range from 0.0038 between *Sintra* and *Monchique* to 0.1225 between *Kenitra* and *Var* (average FST of 0.0553).

**Figure 2:**
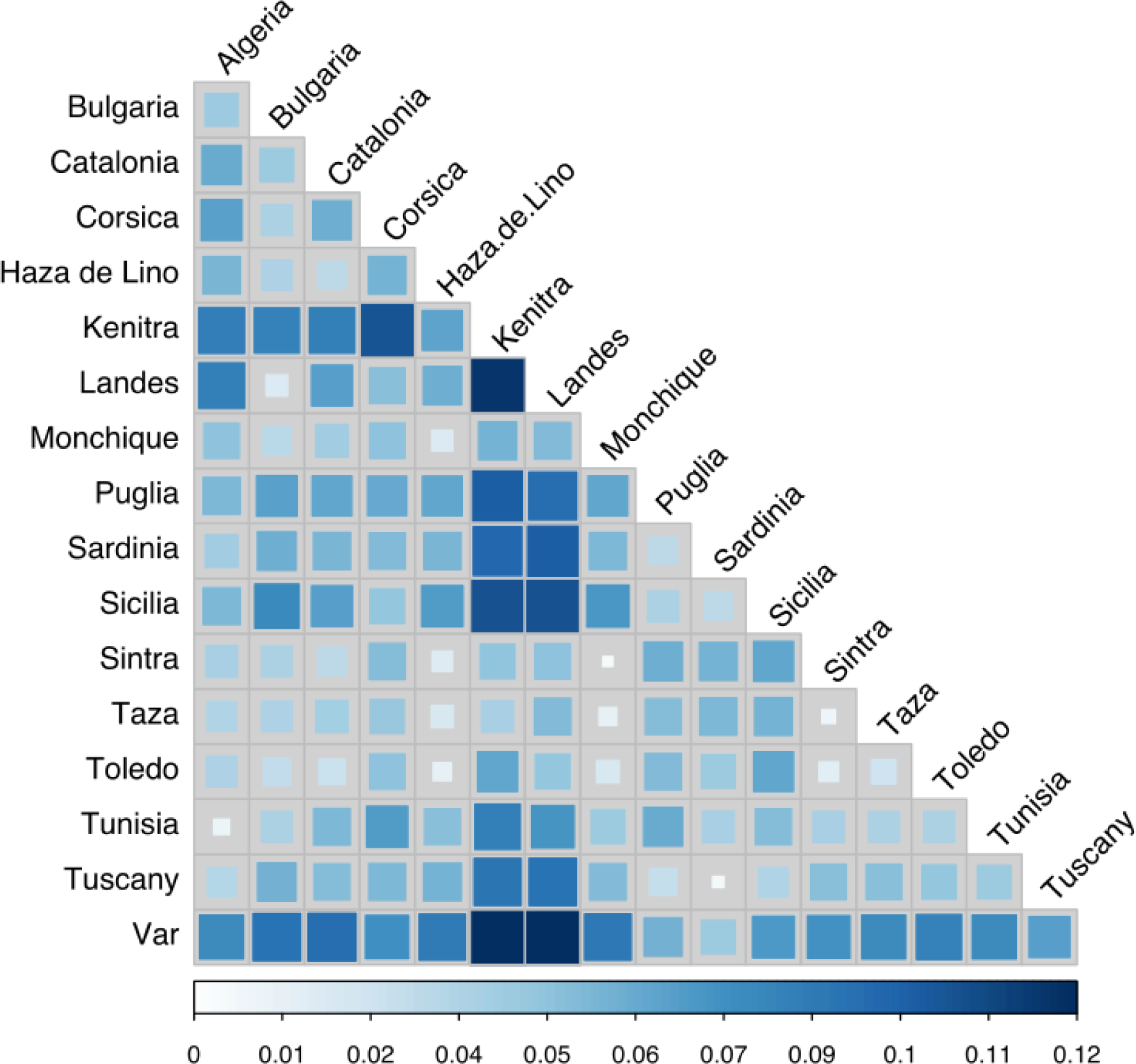
Pairwise Fst plot between sampling sites. Darker blue represents a higher pairwise Fst value, and lighter blue represents a lower value.

Hardy-Weinberg Equilibrium tests revealed that a heterozygote deficit exists in most sampling sites (Table 2), in fact, only *Bulgaria* and *Landes* sampling sites seem not to have an excess of homozygote individuals. When looking at HWE results per marker, of the 2,547 SNPs, only 109 reveal a heterozygote deficit, whereas 23 reveal a deficit of homozygotes. Performing the same test on all individuals as a single large population also revealed a deficit of heterozygotes. The performed Mantel test revealed no evidence of IBD among *Q. suber* individuals.

### 3.1 Outlier detection and environmental association

Population differentiation and ecological association approaches (François, Martins, Caye, & Schoville, 2016) were employed aiming at the identification of loci targeted by selection. In the first strategy, highly differentiated loci among populations, measured as outliers in Fst distribution, were detected by the software *BayeScan* and *SelEstim* uncovering 32 and 48 outlier SNPs respectively (Supplementary Table 5). Most of the loci considered under outliers by *BayeScan* were also present in the set of loci fagged as outlier by *SelEstim.* This set of 31 common markers was considered as being putatively under the effect of natural selection.

For a functional characterization of these loci, the draft genome sequence of *Q. lobata* was used as a proxy for similarity searches. Ten of the 31 sequences revealed significant matches to *Q. lobata* genome scaffolds. Of these, seven were not annotated, and four could be matched to an annotated region (Table 3).

**Table 3:**
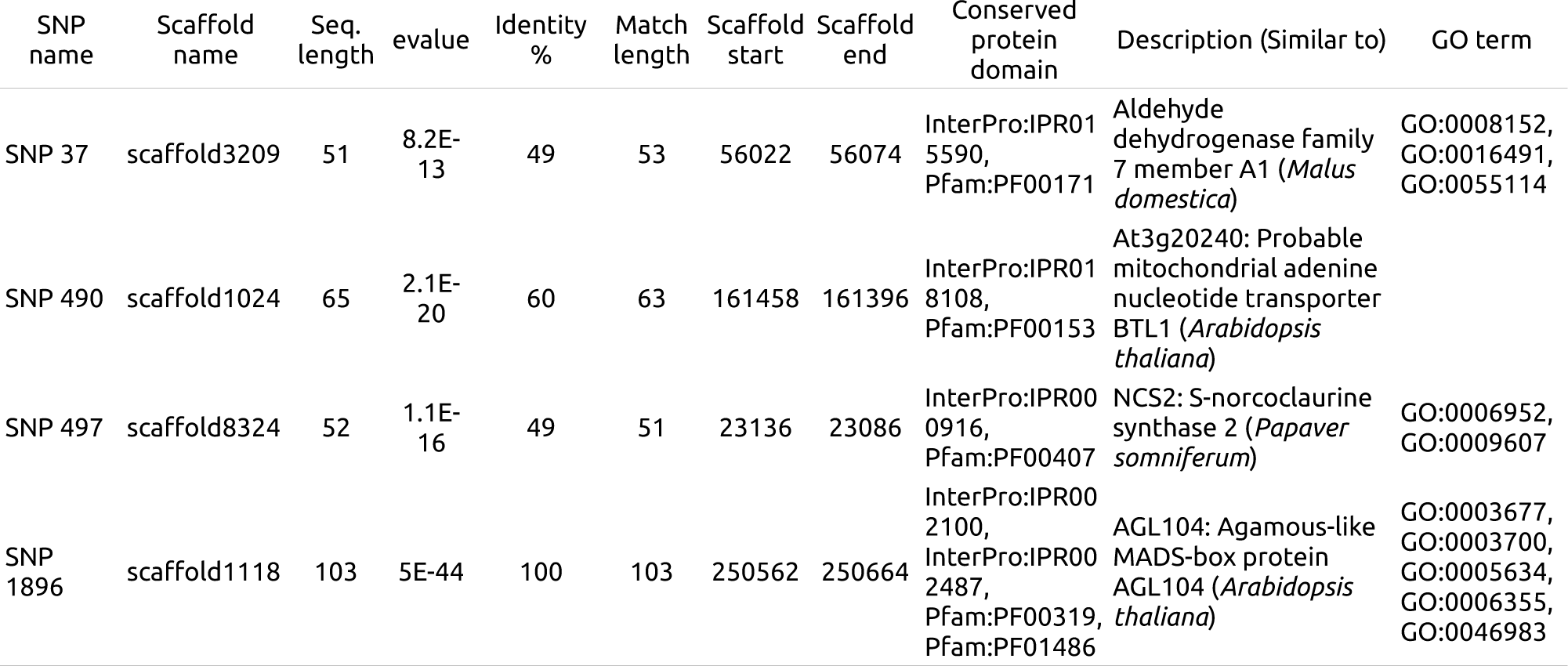
Summary of best BLAST hit results for loci with SNPs considered outliers against the genome of *Q. lobata.*

The ecological association approach was carried out using the software *BayPass* and yielded 374 associations between 329 SNPs and 14 of the 16 tested environmental variables (no associations were found with neither “Temperature Annual Range” nor “Precipitation Seasonality”). These associations can be found in Supplementary Table 6. Despite this relatively high number of associations, it is important to note that 72 of these associations were between a SNP and a geospatial variable: 9 associations with “Latitude”, 55 with “Longitude” and 8 with “Altitude”. Of all environmental variables, the one with most markers associated is “Precipitation of Driest Month” with 79 associations, followed by “Mean Temperature of Driest Quarter” with 51 associations, and “Temperature Seasonality” with 33 associations.

Sequences containing 144 of the 329 markers associated with environmental variables were matched to entries in the *Q. lobata* genome, however, of these only 47 were annotated (Table 4).

**Table 4:**
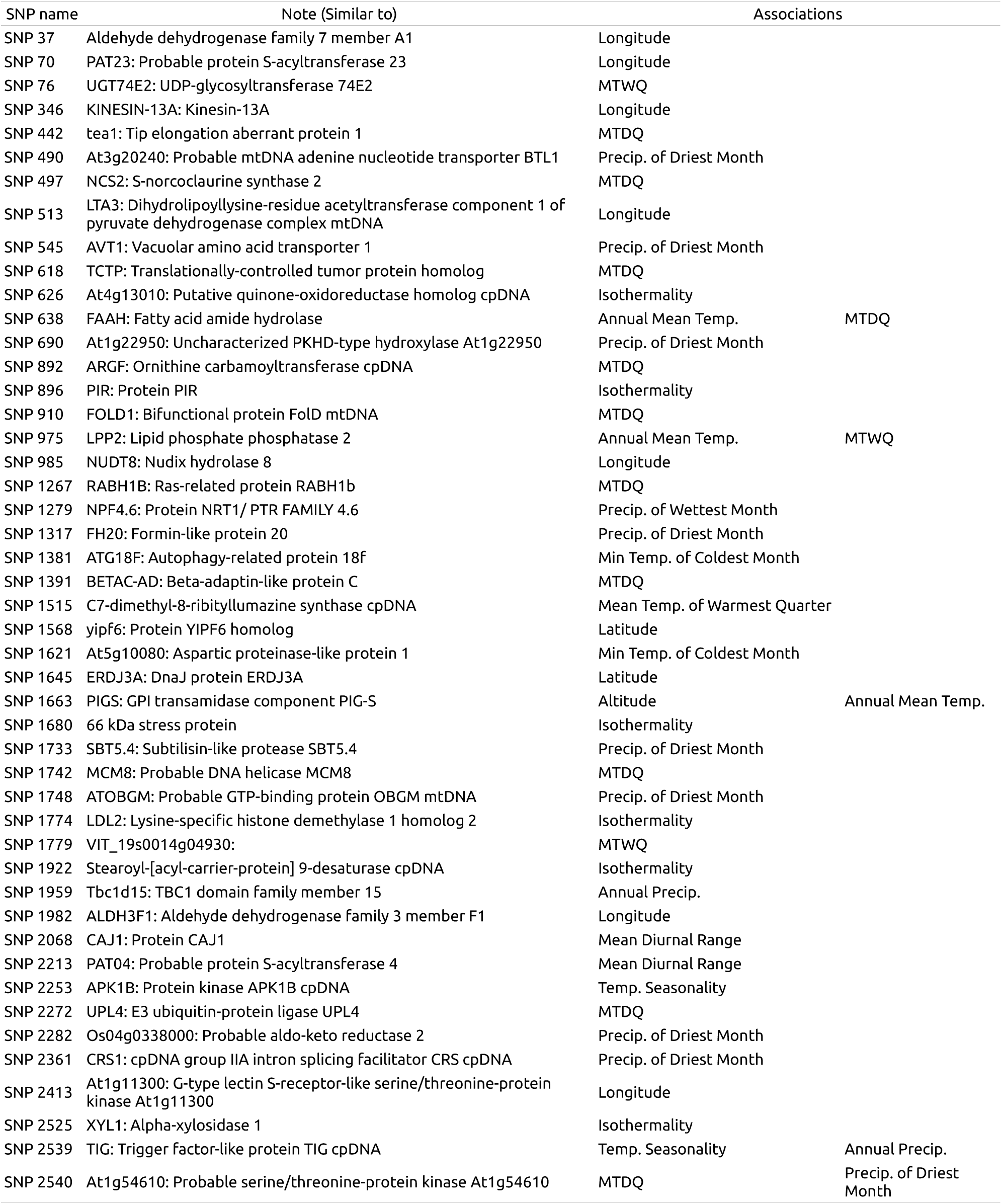
Summary of BLAST hits for loci with SNPs associated to one or more environmental variables. “MTDQ” and “MTWQ” stand for “Mean Temperature of Driest Quarter” and “Mean Temperature of Wettest Quarter” respectively.

The union of the outlier loci set and the set of loci associated with at least one environmental variable resulted in a dataset of 341 SNPs which were deemed “nonneutral” (19 SNPs were common to both loci sets). The remaining 2206 SNPs were grouped in another sub-dataset, deemed “neutral”.

### 3.2 Population structure

Clustering analyses were used to infer the current population structure of *Q. suber* in the West Mediterranean. The *>TI*> method implemented in the software *MavericK* determined the best “K” value to be “1” on all datasets. The classic method for the structure software, the *evanno* method revealed that K=2 had the best ΔΚ, followed by K=3 and K=4 on all datasets. It is, however, important to note that the *evanno* method is not able to evaluate the ΔΚ value for K=1. Despite this assessment, the presented plots are always with K=2, but with strong evidence that that the most likely scenario is that there is no structuring of any kind.

The Q-matrix plot showing the relatedness of each genotype to each considered deme of *MavericKs* results produced using all loci (Figure 3A) can be interpreted as a rough split between Western individuals (from locations *Sintra, Monchique, Kenitra, Toledo, Landes, Taza, Haza de lino* and *Catalonia)*, which are mostly assigned to cluster “1” and Eastern ones (from locations *Var, Algeria, Sardinia, Corsica, Tunisia, Tuscany, Sicilia, Puglia* and *Bulgaria)*, which are mostly assigned to cluster “2”. Individuals from *Bulgaria* are a notable exception, since individual genotypes are mostly assigned to cluster “1” similar to those of individuals from Western locations (due to the species’ introduced origin (Varela, 2000)). However, this West - East split is somewhat fuzzy, as individuals’ genomes are never completely attributed to a single cluster. In fact, most individuals have a considerable part of their genome attributed to both cluster “1” and “2”. Furthermore, individuals from some eastern locations have their genomes mostly attributed to cluster “1” (Var *21, Corsica 3, Corsica 11, Corsica 14* and *Puglia 5)*, and individuals from *Tunisia* are almost equally split between both clusters.

**Figure 3:**
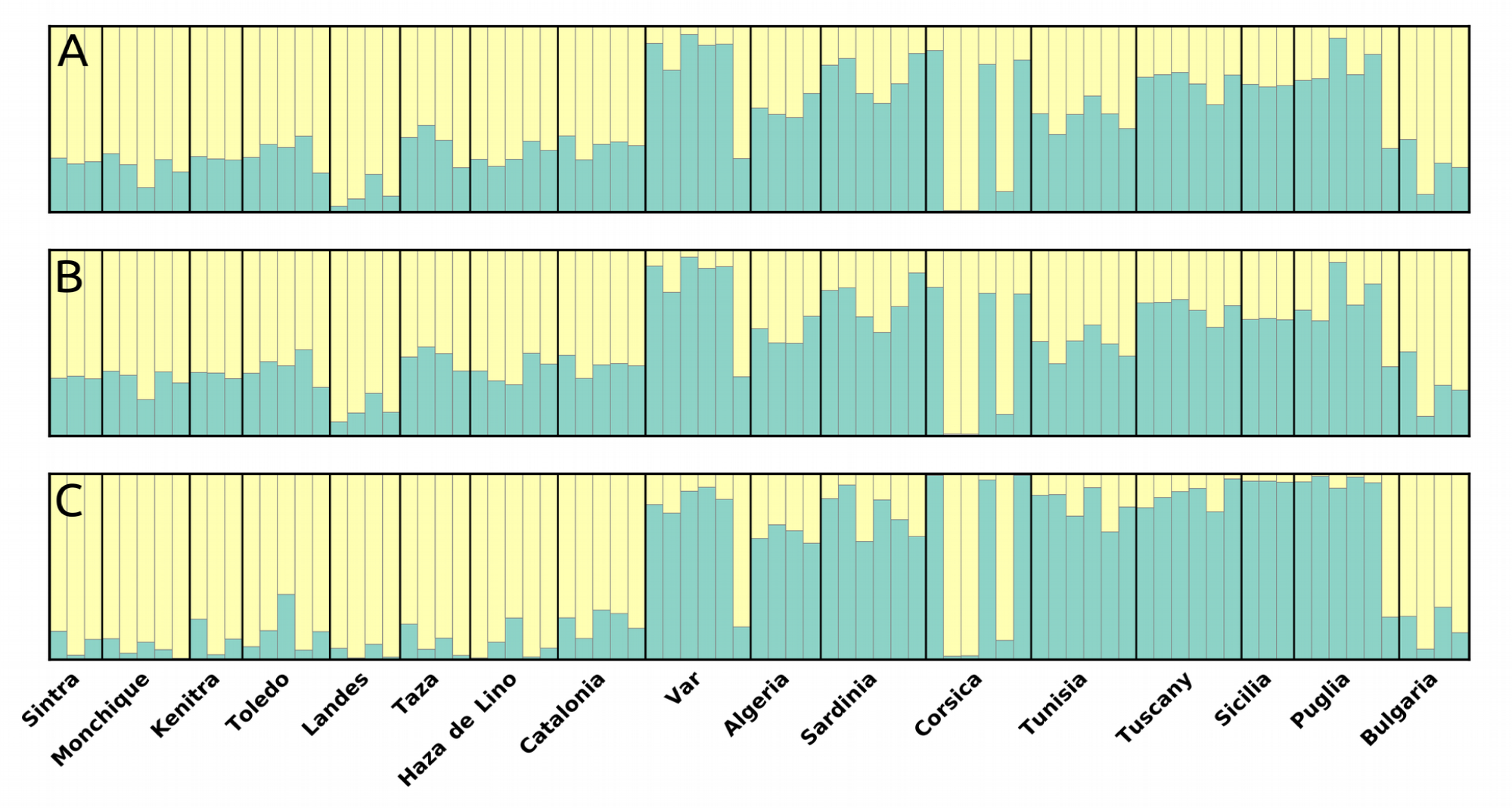
*MavericK* clustering plots for K=2. Sampling sites are presented from West to East. “A” is the Q- value plot for the dataset with all loci, “B” is for the dataset with only “neutral” loci, and “C” if for the dataset with only “non-neutral” loci.

The Q-plot obtained using the “neutral” loci subset (Figure 3B) is nearly identical to the one with all the loci, and can be interpreted in the same way.

The Q-plot produced using only the 305 (13.4%) “non-neutral” loci (Figure 3C), however does bear a different clustering pattern from the previous ones. In this case, the East - West split is more evident, as individual genomes’ attribution to each cluster is not as evenly split, but rather displays a more pronounced individual genome attribution to either cluster.

The Q-plot obtained from structure (Supplementary Figure 1) reveals a similar pattern to that of *MavericK* on all datasets.

The PCA clustering method (largest eigenvector values of 0.0431 and 0.0241) is essentially concordant with the previous methods, revealing two loosely defined groupings (Supplementary Figure 2). The first group containing individuals from *Algeria, Corsica, Puglia, Sardinia, Sicilia, Tuscany, Tunisia* and *Var* and the second group containing individuals from *Bulgaria, Catalonia, Corsica, Haza de Lino, Kenitra, Landes, Monchique, Puglia, Sintra, Taza, Toledo* and Var. The groups are loosely defined, because they somewhat resemble an East - West split, but individuals from *Corsica, Puglia* and *Var* are present in both groups. Just as in the Q-plots, Bulgarian individuals group with Western ones, despite existing on the edge of the species’ Eastern range. Finally, a less pronounced sub-grouping is discernible: one comprising three individuals from *Corsica;* a second comprising all *Landes* individuals, plus three individuals from *Bulgaria;* and a third sub-group consisting of two individuals from *Puglia* and three from *Var*.

### 3.3 Risk of non-adaptedness (RONA)

A summary of the RONA analyses for both a low emission scenario (RCP26) and a high emission scenario (RCP85) predictions can be found in Figure 4 and Supplementary Table 7. The most represented environmental variables are “Precipitation of Driest Month” (79 SNPs, mean R^2^=0.1597), “Mean Temperature of Driest Quarter” (51 SNPs, mean R^2^=0.1466) and “Temperature Seasonality” (33 SNPs, mean R^2^=0.1545). The values of RONA per sampling site are always higher for RCP85 than for RCP26, except for “Precipitation of Driest Month” in *Tunisia* where RCP85 has a lower RONA than RCP26, and in *Kenitra* where they are the same (the “Precipitation of Driest Month” variable in *Kenitra* is not predicted to change from current conditions (0 mm2), regardless of the model).

**Figure 4:**
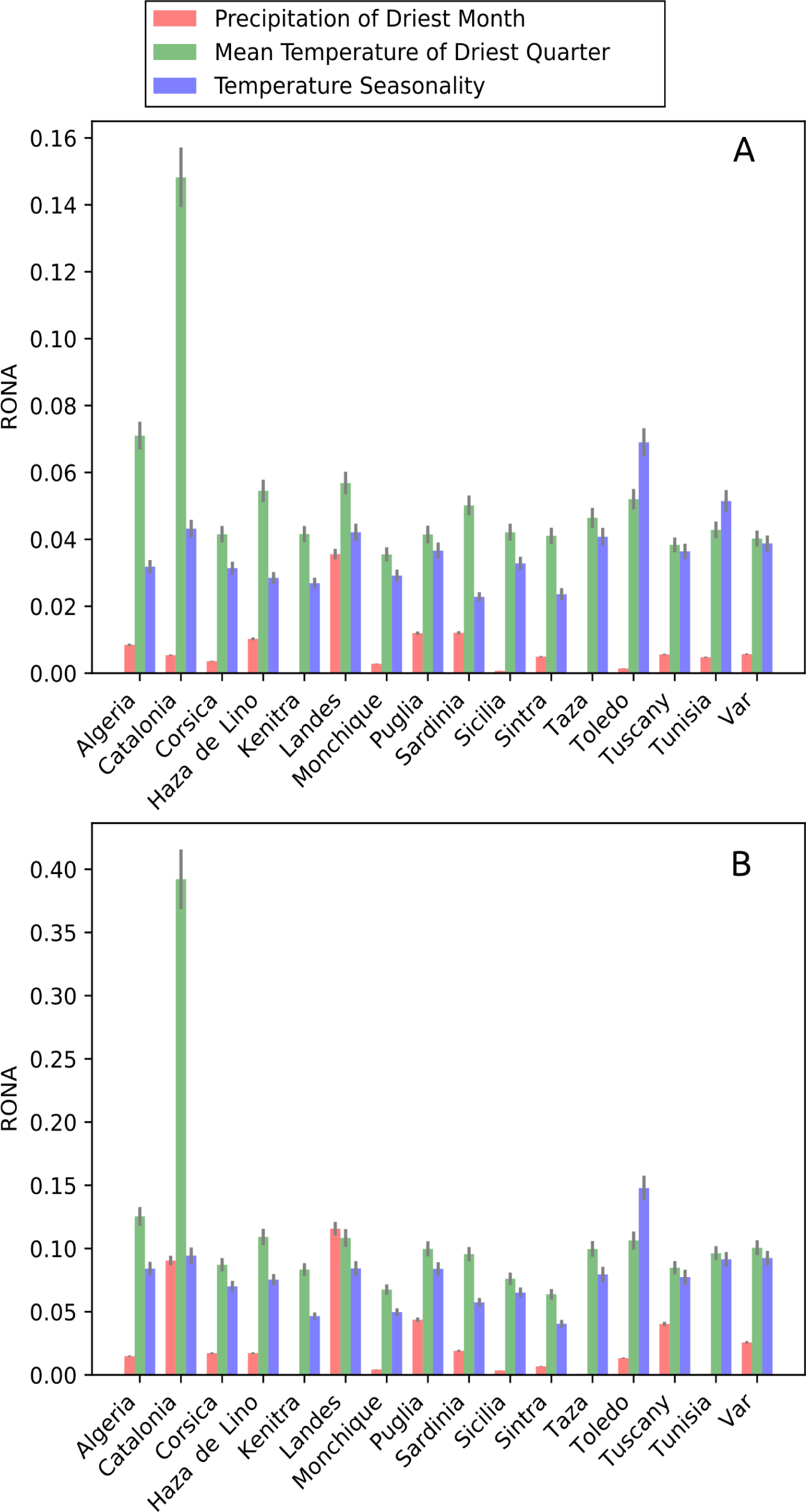
Risk of Non-Adaptedness plot for the three SNPs with most associations. Bars represent weighted means (by R2 value) and lines represent standard error. (A) is the plot for RCP26 and (B) is for RCP85 prediction models.

Under the RCP26 predictions, the highest RONA values for “Mean Temperature of Driest Quarter” is *Landes* (0.1482), for “Temperature Seasonality” is *Toledo* (0.0690) and for “Precipitation of Driest Month” is *Landes* (0.0356). Under the RCP85 predictions, *Catalonia* presents the highest values of RONA for “Mean Temperature of Driest Quarter” (0.3921), *Landes* presents the highest RONA for “Precipitation of Driest Month” (0.1157), whereas *Toledo* has the highest value (0.1478) for “Temperature Seasonality”. It is important to note that the high RONA values of *Catalonia* are twice as high as the second highest RONA value on the RCP26 prediction and more than three times as high for RCP85.

## 4 Discussion

In this study, *Quercus suber* individuals were sampled across the species’ distribution range to assess the population structure, impact of local adaptation and provide an estimate of the RONA value of each sampled location.

Due to the relatively large size of *Q. subefs* genome (Zoldos, Papes, Brown, Panaud, & Siljak-Yakovlev, 1998) a genome reduction technique, GBS, was used to discover SNPs for this species. There is no “standard” parameter set to call SNPs on GBS datasets, since this will ultimately depend on the organism being studied. The stringent approach used in this study was, however, deemed preferable to alternatives that could result in more SNPs being called at the cost of lowering confidence in the called variants, eventually biasing analyses results. In fact, since no biological replicates were performed for this study, a conservative approach was always preferred as to minimize biases in the results.

After stringent quality filtering, a set of 2,547 SNPs was used in this study. This number is lower than that of some studies with similar data (Berthouly-Salazar et al., 2016), which obtained ~22k SNPs (albeit using a more frequent cutting enzyme), but still more than (De Kort et al., 2014), which obtained 1630 SNPs, very close to that of (Escudero, Eaton, Hahn, & Hipp, 2014) and (Pais, Whetten, & Xiang, 2017). Even though this number may seem small, in the universe of *Q. suber*’s genome of ~750 Mbp, this is to date the largest number of molecular markers available for this species and represents a step forward to increase the power of population genetics studies.

### 4.1 Population genetic structure

Past studies (Magri et al., 2007) have characterized *Q. suber* as a highly structured species, with an evolutionary history shaped by large effect events, such as plate tectonics. These were, however, mostly based on plastidial DNA data, which is known to not always provide a comprehensive view on a species’ evolutionary history (Kirk & Freeland, 2011). The nuclear markers developed for this work provide a somewhat different perspective.

The obtained values of F_IS_ are higher than those of unstructured European oaks when analysed with the same type of markers, such as *Quercus robur* or *Quercus petraea* (Guichoux et al., 2013), but are nonetheless relatively low in general, which is compatible with low levels of population structuring.

Only two sampling sites did not reveal significant deviations from HWE *(Bulgaria* and *Landes)* regarding heterozygote deficiency. No sampling site exhibited heterozygote excess. Although this pattern is not usual, few individual markers deviate from HWE (4.28% reveal excess heterozygotes and 0.90% deficit heterozygotes). This may be due to the fact that each sampling site does not represent a real biological population (albeit the same pattern arises when all individuals are merged into a single group, or in Eastern and Western groups), or to non random mating across the species distribution range.

Similar to what is observed with F_IS_, F_ST_ values are on average (0.0553) higher than on unstructured trees species (0.0125) (Guichoux et al., 2013), but lower than other well structured trees such as eucalypts (0.095) (Cappa et al., 2013). This data supports what the clustering analyses reveal: an incomplete segregation in two clusters, as seen on Figure 3. Although clustering analyses using all loci do not provide a clear structuring signal (and the “TI” method clearly favours a scenario of a single large panmictic population), the produced *Q. suber* Q-plots do show some degree of segregation between Western and Eastern individuals, which could hint at some form of local adaptation.

A comparative Q-plot analysis between “neutral” and “non-neutral”, however, reveals the most contrasting differences regarding *Q. subefs* population structure.

In Figure 3C, where the Q-plot was produced using only loci putatively under selection, the division between Western and Eastern individuals is clearer than in Figure 3A and Figure 3B. Conversely, in Figure 3B, which was drawn based on loci deemed “neutral”, a pattern very similar to the Q-plots of all loci emerges, which supports a scenario of an incomplete segregation between individuals from Eastern and Western locations. This evidence, combined with the relatively low pairwise F_IS_ and F_ST_ values, suggests a balance between local adaptation and gene flow. Whereas the former is responsible for maintaining the species’ standing genetic variation across the species range and the latter for the species’s response to local environmental differences. Intense gene flow would also explain the relatively low proportion of outlier SNPs, which may be counteracting reactions to weak selective pressures. At the same time, this balance may provide the species a relatively large genetic variability to respond to strong selection (De Kort et al., 2014; Kremer et al., 2012).

Data from this work does not seem to support the four lineages hypothesis proposed in (Magri et al., 2007). It could be argued that these plastidial lineages exist due to population contractions and expansions from glacial refugia, but a high gene flow would have erased any evidence of their existence in the nuclear genome, as is thought to have occurred in other tree species (Eidesen et al., 2007). However it seems just as likely to assume a scenario without refugia, where *Q. suber* maintained a relatively large population effective, even during glacial periods, and the cpDNA segregation is a consequence of different dispersal capacities of pollen and acorns (Sork, 1984).

Two hypotheses can thus be proposed to explain the currently observed genetic structure. (1) balance between gene flow and local adaptation is responsible for both creating and maintaining the current level of nuclear divergence. Whereas local adaptation tends to cause divergence between contrasting regions, species wide gene flow counters this with an homogenising effect. Population contractions in refugia locations during glacial periods explain both the plastidial lineages and, to some extent, the small difference between Eastern and Western locations. (2) In this scenario, balance between gene flow and adaptation is responsible for maintaining the current genetic pattern, but not for their origin, which is explained by differential hybridization of *Q. suber* with *Q. cerris* in the East (Bagnoli et al., 2016) and with *Q. ilex s.l.* in the West (Burgarella et al., 2009). Combination of the two above phenomena is the cause for the small East-West differentiation. Under this hypothesis, *Q. suber* would maintain a high nuclear population effective, even during glacial periods, but restrict plastidial lineages’ geographic scope, as suggested in (López de Heredia, Carrión, Jiménez, Collada, & Gil, 2007), since *Q. suber* always acts as a pollen donor in these hybridization events (Boavida, Silva, & Feijó, 2001). This would cause a particularly large difference in effective population size between nuDNA and cpDNA, which explains why cpDNA can be divided into lineages - random mutations crop up in different geographic locations and are differentially maintained by drift. The SNP data is compatible with both hypothesis, but not sufficient to confirm any of them, and as such, the issue will remain open for investigation.

### 4.2 Outlier detection and environmental association analyses

The method used to detect outlier loci fagged ~1.2% of the total SNPs, which is in line with what was found on other similar studies (Berdan, Mazzoni, Waurick, Roehr, & Mayer, 2015; Chen et al., 2012). Of the 31 outlier markers found, only four had a match to an annotated location in *Q. lobata*’s genome. This low proportion is likely due to a combination of factors, such as the distance between *Q. suber* and *Q. lobata*, and the incomplete annotation of *Q. lobata’s* genome. On the other hand, it emphasizes the need for more genomic resources in this area, which can potentially provide important functional information of these SNPs in *Q. suber*’s genome, that will at least for now remain unknown. Of particular note is SNP 493, whose sequence is a match to a region of the *Q. lobata* genome, annotated as “Similar to NCS2: S-norcoclaurine synthase 2 *(Papaver somniferum)”*, a protein family member usually expressed upon infections and stressful conditions (van Loon, Rep, & Pieterse, 2006). This can be a particularly interesting marker for downstream studies regarding adaptation to infection response.

The environmental association analyses (EAA) served two purposes in this work. On one hand, the reported associations work as a proxy for detecting local adaptation, and on the other hand, allow the attribution of a RONA score to each sampling site. *Q. suber* is known to be very sensitive to precipitation and temperature conditions (Vessella et al., 2017, and as such, it was expected beforehand that some of the markers obtained in this study were to be associated with some of these conditions (Rellstab et al., 2016). In order to understand how important the found associations are for the local adaptation process, it is necessary to understand the putative function of the genomic region where each SNP was found. Querying the available sequences against *Q. lobata’s* genome annotations, has provided insights regarding some of the markers’ sequences putative function. The proportion of sequences that were a match to an annotated region, however, is rather small - only ~14.3% of the queried sequences could be matched to such regions.

Of the 47 SNPs associated with an environmental variable that returned hits to annotated regions of *Q. lobata’s* genome, four are likely located in a mitochondrial region, seven in chloroplastidial regions, and 36 in nuclear regions. While all these associations are potentially interesting to explore, doing so falls outside the grander scope of this work. Nevertheless, 6 SNPs are particularly interesting to take a closer look at, mostly due to how much information is available regarding the identified genomic region function.

In addition to being identified as an outlier, SNP 497 is also associated with the variable “Mean Temperature of Driest Quarter”. It is interesting to assess that a marker located in a genetic region known to be expressed during stressful conditions is associated with an environmental variable that cork oak is known to be sensitive to. This makes SNP 497 a very interesting candidate for downstream studies.

SNP 638 is located in a sequence annotated as “Similar to FAAH: Fatty acid amide hydrolase”. This is a family of proteins that are known to play a role in the transport of fixed nitrogen from bacteroids to plant cells in symbiotic nitrogen metabolism (Shin et al., 2002). *Q. suber* is known to have symbiotic associations with mycorrhizae (Sebastiana et al., 2014) and the association of this marker with both “Annual Mean Temperature” and “Mean Temperature of Driest Quarter” can lead to important findings on downstream studies.

SNP 1621 and SNP 1733 are located in sequences that matched regions whose annotation indicates they may be involved in pathogen defence signalling (Figueiredo, Monteiro, & Sebastiana, 2014; Xia et al., 2004). The matched annotations are “Similar to At5g10080: Aspartic proteinase-like protein 1” and “Similar to SBT5.4: Subtilisin-like protease SBT5.4” respectively. SNP 1621 is associated with the variable “Min Temperature of Coldest Month”, and SNP 1733 is associated with “Precipitation of driest month”. Like the above, these markers can be potentially very interesting for downstream analyses regarding pathogen response.

SNP 1645 is located in a sequence that matched a region annotated as “Similar to ERDJ3A: DnaJ protein ERDJ3A”. This protein is known to play a role in pollen tube formation during heat stress (Yang et al., 2009). In this case, the maker is associated with “Latitude”, which might be working as a proxy for some temperature related variable that was not used in this study.

The sequence where SNP 2272 is found can be matched to a region annotated as “Similar to UPL4: E3 ubiquitin-protein ligase UPL4”. This family of proteins is known to be involved in leaf senescence processes (Miao & Zentgraf, 2010). Its association with “Mean Temperature of Driest Quarter” makes SNP 2272 a good candidate for downstream research regarding *Q. subefs* leaf development.

### 4.3 Risk of non-adaptedness

Although the RONA method is a greatly simplified model (its limitations are described in (Rellstab et al., 2016)), it provides an initial estimate of how affected *Q. suber* is likely to be by environmental changes (at least as far as the tested variables are concerned). The implementation developed for this work, named *pyRONA* suffers from most of the same limitations as the original application, even though it is based on an arguably superior association detection method (Gautier, 2015), but introduces a correction to the average values based on the R^2^ of each marker association (by using weighted means). The automation brought by this new implementation, easily allows two different emission scenarios (RCP26 and RCP85) to be tested and compared.

With the exception of *Catalonia*, which seems to have an exceptionally high highest RONA value under both prediction models, the other locations present relatively low RONA values for the tested variables. The variable “Mean Temperature of Driest Quarter” appears to be the tested variable that requires the greatest changes in allele frequencies to ensure adaptation of the species to the local projected changes, although “Temperature Seasonality” is not far behind. These RONA values, are nevertheless smaller than those presented in (Rellstab et al., 2016). This might be due to various factors, such as the different variables tested, the geographic scope of the study, the species’ respective tolerance to environmental ranges, the differences between species’ standing genetic variation, the association detection method, or likely a combination of several of these factors.

Notwithstanding, the obtained results seem to indicate that *Q. suber* is generally well genetically equipped to handle climatic change in most of its current distribution (with the notable exception of *Catalonia*). Despite cork oak’s long generation time, it seems reasonable that during the considered time frame current populations are able to shift their allele frequencies (2% to 10% on average, depending on the predictive model) due to the species relatively high standing genetic variation, which according to (Kremer et al., 2012) should really work in the species’ favour in the presence of strong selective pressures.

This study, however, is limited to the considered environmental variables. Other factors that were not included in this work may have a larger effect on *Q. subefs* RONA. Inferring future adaptive potential of species is not yet commonplace practice (Jordan, Hofmann, Dillon, & Prober, 2017; Rellstab et al., 2016), however, combining this type of study with ecological niche modelling approaches has the potential to greatly improve the accuracy of both kinds of predictions.

## 5 Conclusions

In this study, new nuclear markers were developed to shed new light on *Q. subers* evolutionary history, which is important to understand, in order to attempt to predict the species response to future environmental pressures (Kremer et al., 2014).

Despite the relatively large geographic distances involved, the nuclear markers used in this work indicate lesser genetic structuring than previously thought from cpDNA markers, that clearly segregated the species in several well defined demes (Magri et al., 2007). The SNP data from this work can thus be used to propose two new hypotheses to replace the current view of a genetic structure carved by population recessions and expansions from glacial refugia. The observed genetic structure origin and maintenance can be explained either by balance between gene flow and local adaptation, or alternatively, differential hybridization of *Q. suber* with *Q. ilex s.l.* in the West and *Q. cerris* in the East is responsible for the geographic differences, which are then maintained by the mentioned balance between gene flow and local adaptation (albeit more research is required to confirm this second hypothesis).

Despite the genetic structure homogeneity, outlier and association analyses hint at the existence of local adaptation. The RONA analyses suggest that this balance, between local adaptation and gene flow, may be key in the *Q. suber’s* response to climatic change. It is also worth considering that despite the species likely capability to shift its allele frequencies for survival in the short term, the effects of such changes in the long term can be quite unpredictable (Feder, Egan, & Nosil, 2012; Lenormand, 2002), and only very recently have they began being understood (Aguilée, Raoul, Rousset, & Ronce, 2016).

This study starts by providing a new perspective into the population genetics of *Q. suber*, and, based on this data, suggests an initial conjecture on the species’ future, despite the used technique’s limitations. Even though studies regarding *Q. subers* response to climatic change are not new (Correia et al., 2017; Vessella et al., 2017), this is the first work where this response is investigated from an adaptive perspective. One aspect that could thoroughly improve its reliability would be the availability of more genomic resources, especially a thoroughly annotated genome of the species. Such resource would allow the identification of more markers, and assess the reliability of more associations, which would also allow a more refined method for assessing which loci are more likely to be under the effects of selection. Fortunately, such efforts are underway, and further work in this area should benefit from it in the near future.

## 6 Acknowledgements

We would like to thank R. Nunes, A. S. Rodrigues, C. Ribeiro and I. Modesto, for their help during sample collection. Funding was provided by projects SOBREIRO/0036/2009 (under the framework of the Cork Oak ESTs Consortium) and UID/BIA/00329/2013 from Fundação para a Ciência e Tecnología (FCT) - Portugal. F. Pina-Martins was funded by FCT grant SFRH/BD/51411/2011, under the PhD program “Biology and Ecology of Global Changes”, Univ. Aveiro & Univ. Lisbon, Portugal.

